# Genomic islands of heterozygosity maintained across caribou populations despite inbreeding

**DOI:** 10.1101/2020.12.29.424772

**Authors:** Kirsten Solmundson, Jeff Bowman, Paul J. Wilson, Rebecca S. Taylor, Rebekah L. Horn, Sonesinh Keobouasone, Micheline Manseau

**Affiliations:** Environmental & Life Sciences Graduate Program, Trent University, 1600 West Bank Drive, Peterborough, ON, K9L 0G2, Canada; Wildlife Research and Monitoring Section, Ontario Ministry of Natural Resources and Forestry, Trent University, DNA Building, Peterborough, ON K9L 0G2, Canada; Biology Department, Trent University, 1600 West Bank Drive, Peterborough, ON K9J 7B8, Canada; Landscape Science and Technology Division, Environment and Climate Change Canada, 1125 Colonel By Drive, Ottawa, ON K1S 5R1, Canada

**Keywords:** Runs of homozygosity, inbreeding, islands of heterozygosity, balancing selection, conservation genomics, caribou

## Abstract

Small, isolated populations are prone to inbreeding, increasing the proportion of homozygous sites across the genome that can be quantified as runs of homozygosity (ROH). Caribou *(Rangifer tarandus)* are declining across their range in Canada; thus, understanding the effects of inbreeding on genetic potential is pertinent for conserving small, isolated populations. We quantified ROH in high-coverage whole genomes of boreal caribou from small, isolated populations in southern Ontario, Canada, in comparison to caribou from the continuous range of Ontario, other caribou ecotypes in Canada, and western Greenland. Sampled populations presented divergent evolutionary histories, differing population sizes, and extents of isolation. We conducted BLAST searches across regions of elevated heterozygosity to identify genes that have maintained variation despite inbreeding. We found caribou from recently isolated populations in Ontario had a large proportion of their genome in long ROH. We observed even larger proportions but shorter ROH in western Greenland, indicating that inbreeding has occurred over a longer period in comparison to other populations. We observed the least inbreeding in barren-ground and eastern migratory caribou, which occur in larger population sizes than boreal caribou. Despite vastly different inbreeding extents, we found regions of high heterozygosity maintained across all populations. Within these islands of heterozygosity, we identified genes associated with immunity, signaling regulation, nucleotide binding, toxin elimination, and feeding behaviour regulation. In this study, we confirm inbreeding in isolated populations of a species at risk, but also uncover high variation in some genes maintained across divergent populations despite inbreeding, suggesting strong balancing selection.

## 1 INTRODUCTION

Small and isolated populations have limited mate choice, which increases the likelihood of inbreeding (Herfindal et al., 2014). One consequence of inbreeding is increased genome-wide homozygosity, which can be quantified as the proportion of the genome in runs of homozygosity (ROH; Szpiech et al., 2013). ROH measure the genomic level of inbreeding without making assumptions about the founders of the populations, and therefore can provide a more accurate estimate of inbreeding than traditional methods, such as a pedigree (Kardos, Luikart, & Allendorf, 2015). ROH have been used to study the consequences of persisting in small and isolated populations, such as the loss of genetic diversity and increased inbreeding that preceded the extinction of an island population of woolly mammoth *(Mammuthus primigenius;* Palkopoulou et al., 2015). More recently, ROH have been used to investigate inbreeding in species of conservation concern. A study of a Scandinavian wolf *(Canis lupus)* population revealed stretches of ROH throughout the genome of wolves born in an isolated population, whereas in immigrant wolves ROH were rare or absent (Kardos et al., 2018). Knowledge of ROH that are shared, or identical by descent, between individuals or populations is vital for designing mitigation plans and identifying potential candidates for translocations for at-risk species, as demonstrated by a recent study of isolated puma *(Felis concolor)* populations (Saremi et al., 2019).

Inbreeding increases the probability that an individual will receive alleles that are identical by descent (IBD), meaning the individual receives the same allele from both parents at a particular locus, resulting in increased genome-wide homozygosity (Kardos et al., 2015). This increased homozygosity can result in reduced survival or reproduction, known as inbreeding depression (Hedrick & Garcia-Dorado, 2016). Although rarely studied in wild populations, inbreeding depression is well documented in captivity; for instance, in captive bred prairie-chickens *(Tympanuchus cupido attwateri)* mortality was positively correlated with both parental relatedness and the genetic inbreeding coefficient (Hammerly, Morrow, & Johnson, 2013). Inbreeding depression is caused by two genetic effects: the increased expression of recessive deleterious alleles, and increased homozygosity at loci with heterozygote advantage (Charlesworth & Willis, 2009). Deleterious alleles are most likely to occur within long ROH, suggesting recent inbreeding enables rare deleterious variants to exist in homozygous form, resulting in inbreeding depression (Szpiech et al., 2013). The inbreeding load of a population is fueled by the appearance of recessive deleterious variants, but these alleles can be purged by purifying selection if given sufficient time, tempering the effects of inbreeding depression (Hedrick & Garcia-Dorado, 2016). Under intensive inbreeding, islands of high heterozygosity within long homozygous stretches can indicate regions that might harbour recessive deleterious alleles or be associated with heterozygote advantage under balancing selection. A study of endangered brown bears *(Ursus arctos marsicanus)* demonstrated fixation by drift of several deleterious alleles; however, high-variation was maintained in regions related to the immune system, olfactory signaling pathways, and digestion despite inbreeding, suggesting balancing selection prevents the loss of variation at important genes (Benazzo et al., 2017).

The caribou *(Rangifer tarandus)* is an iconic species in Canada that has experienced dramatic declines in both range and population size over the past century, raising conservation concerns (Festa-Bianchet, Ray, Boutin, Côté, & Gunn, 2011; Laliberte & Ripple, 2004). Caribou are a religious, cultural, and social symbol to many Indigenous people in Canada, as well as an important food source in some communities (Festa-Bianchet et al., 2011; Polfus et al., 2016). Caribou diversity is described by different subspecies and ecotypes, which differ in morphology and behaviour; for example, barren-ground caribou *(R. t. groenlandicus)* congregate in large, migratory groups on the tundra (COSEWIC, 2016). Conversely, the woodland subspecies (*R. t. caribou*) has several ecotypes associated with different habitats such as caribou found in the mountains across western Canada (COSEWIC, 2014b), the eastern migratory caribou that migrate between the boreal forest and the tundra in eastern Canada (COSEWIC, 2017a), and boreal caribou that are more sedentary and found throughout the boreal forest (COSEWIC, 2014a). The diversity found in caribou has resulted in the classification of 12 Designatable Units by the Committee on the Status of Endangered Wildlife in Canada (COSEWIC, 2011). Despite this diversity, all caribou in Canada are currently listed as at risk of extinction (Special Concern, Threatened, or Endangered) by COSEWIC (COSEWIC, 2014-2017).

Recent declines in caribou ranges and population sizes have resulted in small and isolated populations, particularly within the sedentary boreal ecotype (COSEWIC, 2014). In Ontario, the range of boreal caribou has been contracting northward for over a century, resulting in isolated populations that have managed to persist along the coast and nearshore islands of Lake Superior, over 150 km south of the continuous range edge (Ontario Ministry of Natural Resources, 2009; Schaefer, 2003). Here, we analyze inbreeding in boreal caribou from Ontario where the range has recently receded, in comparison to boreal caribou from the continuous range in Ontario and

Manitoba, and other caribou ecotypes in central and eastern Canada, and western Greenland. We included eastern migratory caribou from populations in Quebec and Ontario that have experienced historic and recent declines (COSEWIC, 2017a), as well as barren-ground caribou from a large population that has not experienced dramatic historic or recent declines (COSEWIC, 2016). We also included two individuals from western Greenland, where populations have declined by up to 90% in the past two decades (Jepsen, Siegismund, & Fredholm, 2002) despite absence of major predators (Cuyler & Østergaard, 2005). Previous research has indicated high levels of inbreeding in the Greenland population we sampled (Jepsen et al., 2002; Taylor et al., 2020).

Rapid declines in caribou range and population sizes have raised conservation concerns, as small and isolated populations are more prone to inbreeding and eventually may fall into an “extinction vortex” and become extirpated (Gagnon, Yannic, Perrier, & Côté, 2019; Gilpin & Soule, 1986). Yet, it remains unclear how inbreeding is affecting caribou, and whether inbreeding depression is a pertinent threat. A recent study correlated heterozygosity and fitness in two rapidly declining eastern migratory caribou populations, and found no evidence of inbreeding depression (Gagnon et al., 2019). However, the two eastern migratory populations studied by Gagnon et al. (2019) have historically experienced dramatic fluctuations in population size (COSEWIC, 2017a), which may have allowed for the purging of recessive deleterious alleles. The effects of inbreeding depression can be resisted by selection preventing the unmasking of deleterious recessive alleles or maintaining heterozygote advantage (Hedrick & Garcia-Dorado, 2016). Balancing selection, specifically, negative-frequency dependent selection has recently been associated with maintaining phenotypic polymorphisms in caribou along an environmental gradient in western Canada (Cavedon et al. 2019), but has not been linked to inbreeding.

We used whole genome sequences to investigate inbreeding in small and isolated populations of boreal caribou from southern Ontario, boreal caribou populations from the continuous caribou range of Ontario and Manitoba, eastern migratory caribou, barren-ground caribou, and caribou from western Greenland (Figure 1). We sampled caribou from populations that differed in evolutionary history, demographic history and extent of isolation. We predicted to find a large proportion of genomic ROH in boreal caribou from the southern range of Ontario (Figure 1), where recent range contraction has resulted in small and isolated populations (Drake et al., 2018; Schaefer, 2003). We expected to detect lower levels of inbreeding, quantified as ROH, in boreal caribou from the continuous range of Ontario and Manitoba, as well as in the eastern migratory caribou from Ontario and Quebec; populations that have experienced recent declines but are not as small and isolated as the southern range of Ontario (COSEWIC, 2014a, 2017a). Further, we predicted barren-ground caribou from the Qamanirijuaq population ranging over northern Manitoba and Nunavut (Figure 1) will have the lowest proportion of their genome in ROH, as they occur in large populations that have not experienced dramatic historical or recent declines (COSEWIC, 2016). Continuous stretches of ROH are broken up by the recombination of DNA through successive mating events (Ceballos, Joshi, Clark, Ramsay, & Wilson, 2018); thus, we expected to find the longest ROH in boreal caribou from the southern range of Ontario, reflecting recent inbreeding caused by anthropogenic range contraction (Schaefer, 2003). Caribou from western Greenland have likely experienced inbreeding over a longer period of time (Jepsen et al., 2002); thus, we predicted a large proportion of their genome will be in short ROH. Finally, we predicted to find genes associated with deleterious recessive alleles or heterozygote advantage in regions where balancing selection has maintained islands of heterozygosity despite inbreeding (Benazzo et al., 2017).

**FIGURE 1.**
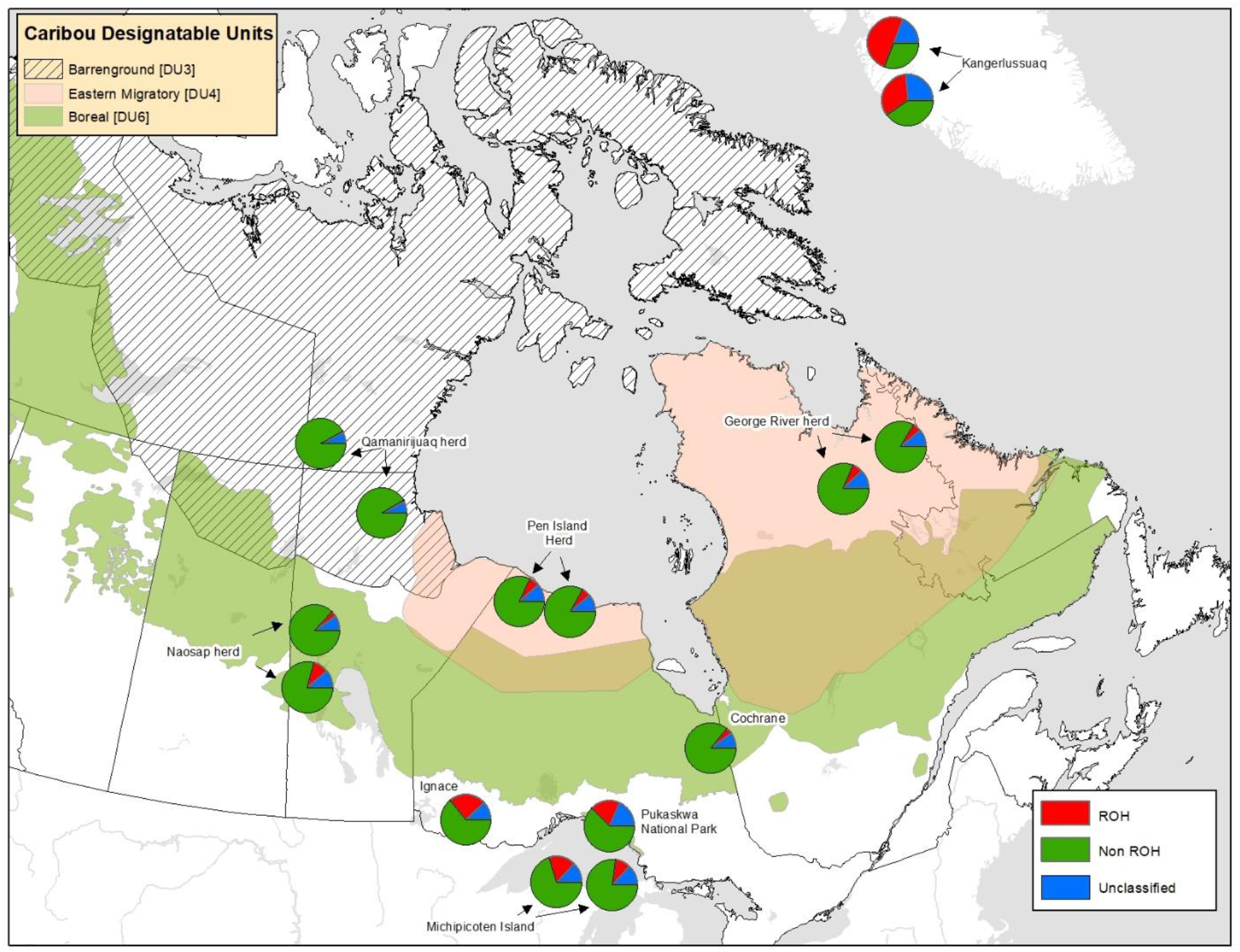
Sample sites of caribou in this study. Background colours show the ranges of the three Canadian designatable units included: barren-ground, eastern migratory, and boreal. Labels on map reflect caribou population names. Pie charts represent the genomic level of inbreeding as the proportion of the genome in ROH, non-ROH, or unclassified for each individual (N=15), at the location that each sample was collected.

## 2 METHODS

### 2.1 Caribou sampling

Tissue samples of three boreal caribou were collected from the southern caribou range of Ontario (two samples from Michipicoten Island, one from Pukaskwa National Park) by provincial biologists and sequenced for the study. Whole-genome raw reads from the continuous caribou range of Ontario, Manitoba, Quebec, and Greenland (Taylor et al. 2020) were used for the remaining populations and can be retrieved from the National Centre for Biotechnology (NCBI) under the BioProject Accession no. PRJNA 634908.

We included four samples from south of the continuous boreal caribou range in Ontario: two samples from the Michipicoten Island and one from the coastal range of Lake Superior (Pukaskwa National Park), as well as one individual from where the continuous caribou range has more recently contracted near Ignace, Ontario (Figure 1). The caribou population on Michipicoten Island was established in the 1980s, when a single bull was sighted on the island and eight additional caribou were translocated from the Slate Islands (Bergerud, Dalton, Butler, Camps, & Ferguson, 2007). The Michipicoten Island population steadily grew, and was estimated to contain 680 caribou in 2011 (Kuchta, 2012); however an especially cold winter in 2014 resulted in ice corridors between the mainland and islands, allowing wolves to colonize the island (Ontario Ministry of Natural Resources, 2018). The caribou population rapidly declined under this new predation pressure, prompting the Government of Ontario to work with partners, including Michipicoten First Nation, to translocate some of the few remaining caribou on Michipicoten Island to other Lake Superior islands: the Slate Islands and Caribou Island (Ontario Ministry of Natural Resources, 2018). Over the past four decades, the coastal caribou population in Pukaskwa National Park has steadily declined and become increasingly isolated from the continuous caribou range of Ontario (Patterson et al., 2014). Caribou disappeared from the park in 2011, and reappeared in 2015, perhaps due to colonization from one of the Lake Superior islands. One of the last caribou captured on wildlife cameras deployed in Pukaskwa National Park had small, malformed antlers, which was suggested to be a sign of inbreeding (Drake et al., 2018). Currently, there are no caribou in the park and the population is considered potentially extirpated.

We also selected one sample from Cochrane, Ontario, which falls along the southern edge of the continuous boreal caribou range and two samples from the Naosap population in Manitoba, which falls within the continuous range and in 2012 was estimated to contain 100-200 caribou (COSEWIC, 2014a). We selected two barren-ground caribou samples from the Qamanirijuaq population that ranges over northern Manitoba and Nunavut, a large population that was estimated to contain 264,661 individuals in 2014 and has not experienced dramatic historical or recent declines (COSEWIC, 2016). Within the eastern migratory ecotype, we included two samples from the George River population, Quebec, which has experienced a dramatic population decline over recent decades from approximately 823,000 individuals in 1993 (Couturier, Courtois, Crépeau, Rivest, & Luttich, 1996), to approximately 8,900 individuals in 2016 (Gagnon et al., 2019); our samples were obtained in 2008, after population declines had already occurred. We also included two eastern migratory caribou from the Pen Island population in northern Ontario, which was estimated to contain 16,638 individuals in 2011 (COSEWIC, 2017a). The eastern migratory caribou populations in Ontario and Quebec are geographically isolated from each other (Figure 1) and recent research has revealed a divergent evolutionary history between these two populations (Taylor et al., 2020). Finally, we selected two samples from the Kangerlussuaq area of western Greenland, where populations have declined by up to 90% in the past two decades (Jepsen et al., 2002). Caribou from this region are geographically separated by the Maniitsoq glacier and have not hybridized with semi-domestic reindeer, unlike some caribou from other regions in Greenland (Jepsen et al. 2002).

### 2.2 Genome sequencing, assembly, and quality control

DNA was extracted from the tissue samples using the Qiagen DNeasy kit and following manufacturer’s protocols (Qiagen, Hilden, Germany). The extracted DNA was then quantified using a Qubit system (Thermo Fisher Scientific, MA, USA) to ensure all samples were above the minimum threshold required for next-generation sequencing (20ng/μL). The extracted DNA was then sent to The Centre for Applied Genomics (TCAG), at The Hospital for Sick Children (Toronto, ON). An Illumina library prep kit (Illumina, San Diego, CA, USA) was used to fragment the DNA and apply sequencing adapters. Each sample was sequenced on one lane of the Illumina HiSeqX platform, yielding paired-end 150bp sequence reads. The raw reads of most samples (N=12) are already available at the National Centre for Biotechnology (NCBI) under the BioProject Accession no. PRJNA 634908. The raw reads of the remaining individuals (N=3) will be made available by time of publication.

We conducted all bioinformatic analyses using cloud computing resources from Compute Canada (RRG gme-665-ab) and Amazon Web Services (https://aws.amazon.com/). First, we removed sequencing adapters and low-quality bases (phred score <30) from the samples using CutAdapt (Martin, 2011) in the program TrimGalore 0.4.2 (Krueger, 2012). We mapped the sequenced reads to the caribou reference genome (Taylor et al., 2019) using Bowtie2 2.3.0 (Langmead & Salzberg, 2012). We used Samtools 1.5 (Li et al., 2009) to convert the SAM file to a BAM file. We then removed duplicate reads and added read group information to each BAM file with Picard 2.17.3 (Broad Institute, n.d.). We sorted the BAM file with Samtools 1.5 and built an index with Picard 2.17.3. Finally, we checked the quality of each BAM file using FastQC 0.11.8 (Andrews, 2010) and calculated the average depth of coverage in ROHan (Renaud, Hanghoj, Korneliussen, Willerslev, & Orland, 2019). All samples (N=15) passed the FASTQC quality assessments and had a high depth of coverage (28-36x; Table 1).

**Table 1.** Information for each caribou included in this study: sampling locations and individual reference number, subspecies classification and Canadian designatable units, population size and extent of isolation, mean depth of coverage from whole genome mapped BAM files, and inbreeding estimates.

| Location and Reference Number | Subspecies | Designatable Unit | Population Size | Extent of Isolation | Mean Depth | Inbreeding Coefficient (F) | F<sub>ROH</sub> | Mean Length ROH |
| --- | --- | --- | --- | --- | --- | --- | --- | --- |
| Ontario Michipicoten Island 39650 | *R. t. caribou* | Boreal | Small | High | 33.6775 | 0.2167 | 0.1664 | 453134 |
| Ontario Michipicoten Island 39651 | *R. t. caribou* | Boreal | Small | High | 35.9172 | 0.2092 | 0.1000 | 301072 |
| Ontario Pukaskwa National Park 39653 | *R. t. caribou* | Boreal | Small | High | 36.3265 | 0.3897 | 0.1902 | 276247 |
| Ontario Cochrane 39654 | *R. t. caribou* | Boreal | Small | Low | 36.4583 | 0.0656 | 0.0403 | 173556 |
| Ontario Ignace 39590 | *R. t. caribou* | Boreal | Small | Moderate | 28.0951 | 0.2398 | 0.2310 | 556507 |
| Manitoba (The Pas) Naosap herd 35324 | *R. t. caribou* | Boreal | Small | Low | 30.3958 | 0.0463 | 0.0963 | 255161 |
| Manitoba (Snow Lake) Naosap herd 35326 | *R. t. caribou* | Boreal | Small | Low | 34.9286 | 0.0358 | 0.0367 | 152613 |
| Manitoba<br>Qamanirijuaq<br>herd 21332 | *R. t.*<br>*groenlandicus* | Barren-ground | Large | Low | 31.8765 | -0.0833 | 0.0123 | 164607 |
| Manitoba<br>Qamanirijuaq<br>herd 21350 | *R. t.*<br>*groenlandicus* | Barren-ground | Large | Low | 30.7167 | -0.0868 | 0.0137 | 152570 |
| Quebec<br>George River<br>herd 27689 | *R. t. caribou* | Eastern<br>migratory | Medium | Low | 31.6268 | 0.0576 | 0.0689 | 208165 |
| Quebec<br>George River<br>herd 27694 | *R. t. caribou* | Eastern<br>migratory | Medium | Low | 32.3503 | 0.0330 | 0.0534 | 151553 |
| Ontario Pen<br>Island herd<br>20917 | *R. t. caribou* | Eastern<br>migratory | Medium | Low | 29.4896 | 0.0062 | 0.0598 | 151538 |
| Ontario Pen<br>Island herd<br>34590 | *R. t. caribou* | Eastern<br>migratory | Medium | Low | 31.5274 | 0.0276 | 0.0679 | 211619 |
| Western<br>Greenland<br>Kangerlussuaq<br>41660 | *R. t.*<br>*groenlandicus* | n/a | Small | High | 33.1561 | 0.5800 | 0.3373 | 308640 |
| Western<br>Greenland<br>Kangerlussuaq<br>41667 | *R. t.*<br>*groenlandicus* | n/a | Small | High | 30.3475 | 0.5877 | 0.5017 | 433852 |

We used the GATK 3.8 (McKenna et al., 2010) Haplotype Caller to produce a variant call format (VCF) file for each caribou. We further used GATK 3.8 to combine and genotype the GVCFs, producing one joint VCF for all samples. We then used VCFtools 0.1.14 (Danecek et al., 2011) to perform additional filtering. We removed indels, sites with a depth <10 or >80, and low-quality genotype calls (score <20). We also filtered to remove genotypes with more than 10% missing data. We did not filter to remove any SNP with a minor allele frequency (MAF) of less than 0.05 as we have only one or two individuals from each location, and thus did not want to remove private sites. The combined VCF file with all caribou contained 28 246 751 SNPs.

### 2.3 Inbreeding analyses

With the combined VCF file, we used VCFtools 0.1.14 (Danecek et al., 2011) to calculate the inbreeding coefficients (F) and relatedness (φ) based on the KING inference (Manichaikul et al., 2010) for all individuals. We also used VCFtools 0.1.14 to calculate a grouped transition/transversion ratio.

We identified runs of homozygosity (ROH) and calculated global heterozygosity rate (θ) from the individual BAM files using the program ROHan (Renaud et al., 2019). ROHan uses a Bayesian approach to identify ROH with a Hidden Markov Model (HMM). For all ROHan analyses we used a sliding window size of 50kb, allowed a maximum heterozygosity level of 0.0005 within ROH, and specified a transition/transversion ratio of 1.97428, based on our calculation from a VCF file that contained all individuals.

ROHan produced a file of heterozygosity estimates for every 50kb window across the genome, as well as a file with the location and length of each ROH. We then plotted the heterozygosity estimates and ROHs across scaffolds of interest to compare patterns between individuals using ggplot2 in R (Wickham, 2016). Namely, from the 40 largest scaffolds, representing approximately 43% of the caribou reference genome, we selected scaffolds where the lower local heterozygosity estimate (θ) exceeded 0.02 (Table S2). We excluded elevated heterozygosity estimates that were located on the edges of scaffolds, as they may be due to sequence error. Within populations where inbreeding was detected (F_ROH_ > 0.1000), we also calculated how many ROH are identical by descent (IBD) or unique using the intersect function in BEDtools (Quinlan & Hall, 2010).

We then conducted BLAST searches across regions of elevated heterozygosity to identify genes that have maintained heterozygosity despite inbreeding. We used NCBI’s nucleotide MegaBLAST algorithm (Agarwala et al., 2016) and masked for lower case letters, as our reference genome contains both hard and soft masking (Taylor et al., 2019). All other BLAST settings were default.

## 3| RESULTS

### 3.1 Inbreeding estimates

The inbreeding coefficient (F) calculated in VCFtools for each individual from the grouped VCF file was highly correlated with genome-wide heterozygosity (θ) and the proportion of the genome in ROH (F_ROH_), which were both calculated in ROHan (Renaud et al., 2019) from individual BAM files (Figure S2).

The highest FROH values were observed in the two caribou from western Greenland (FROH = 0.3373, 0.5017), with ROH comprising half of the genome of one of the individuals (Figure 1). FROH values were also elevated in boreal caribou from Ignace, Ontario that have recently become isolated from the continuous caribou range due to northward range contraction (F_ROH_ = 0.2398), as well as in boreal caribou from Lake Superior (Pukaskwa National Park and Michipicoten Island) that are located over 150 km south of the continuous range edge (F_ROH_ = 0.1000 - 0.1902; Figure 1). The remaining individuals had F_ROH_ values between 0.0123 and 0.0963, with the two notably lowest values corresponding to barren-ground caribou (Table 1).

We investigated the average length of ROH across each genome to estimate whether inbreeding occurred recently or historically; the two individuals with the longest average ROH were observed in populations located south of the continuous caribou range in Ontario (Ignace and Michipicoten Island). Caribou from western Greenland also had long ROH, although they were shorter than predicted based on FROH (Figure 2).

**FIGURE 2.**
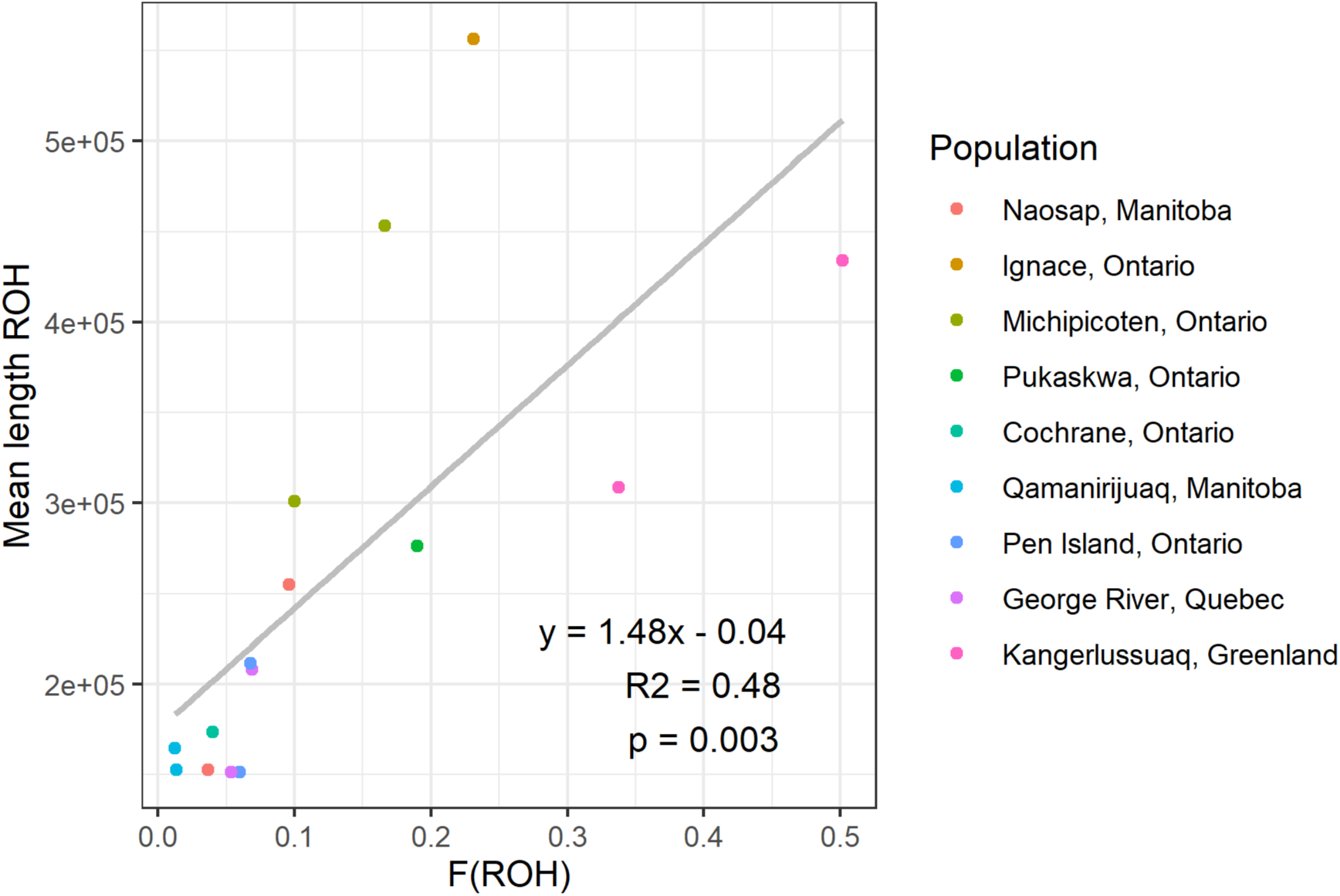
Proportion of the genome in ROH (F_ROH_) versus average length of ROH for each caribou based on high coverage whole genome sequences. Individuals are coloured by the population they were sampled from. The equation describes the line of best fit and R^2^ is the adjusted R-squared value.

### 3.2 Identical by descent ROH

We calculated relatedness between all individuals (φ; Table S1), as well as how many ROH were identical by descent (IBD) for populations where F_ROH_ was greater than 0.10 (Table 1) to gain further insights into their inbreeding histories.

The two boreal caribou *(R. t. caribou)* from Michipicoten Island, Ontario shared 333 of their ROH, corresponding to 38% IBD ROH. The two Michipicoten Island caribou were related to each other (φ = 0.08) but were not related to any other population examined, based on negative relatedness values (Table S1). We then compared other Ontario boreal caribou to determine how many segments were shared with either individual from Michipicoten Island. A caribou from a nearby mainland in Pukaskwa National Park, Ontario shared 62% (N=1028) of its ROH with both individuals from Michipicoten Island but was not related to the Michipicoten caribou (Table S1). The boreal caribou with the highest inbreeding estimate (F_ROH_ = 0.23) was from Ignace, Ontario, and shared 83% (N=713) of ROH segments with the Michipicoten Island caribou and also showed no evidence of relatedness to the island caribou (Table S1). Out of the Ontario boreal caribou sampled, an individual from Cochrane, Ontario shared the lowest amount (N=329) of ROH segments with Michipicoten Island, corresponding to 59% ROH that were IBD, and showed no evidence of relatedness to Michipicoten caribou (Table S1). The caribou from Cochrane also had undergone less inbreeding than the other Ontario boreal caribou sampled (Table 1).

Caribou from western Greenland had the highest counts of shared ROH (N=2406), corresponding to a proportion of 92% of ROH segments that were IBD. However, the relatedness estimate (φ) between these two individuals was 0.09, corresponding to 2^nd^ degree (φ = 0.125) or 3^rd^ degree relatives (φ = 0.0625), which would increase the proportion of the genome that is IBD. The two caribou from western Greenland shared low relatedness values with all of the Canadian caribou (Table S1).

### 3.3 Islands of heterozygosity

We found multiple regions across the genome where high heterozygosity was maintained across all populations, even in individuals that have experienced high levels of inbreeding. In the 40 largest scaffolds, which represented approximately 43% of the caribou genome, we found 17 scaffolds that had a peak of heterozygosity exceeding 0.02 (Table S2). Caribou from western Greenland had many segments of ROH, and boreal caribou from the southern discontinuous range of Ontario had notably long, continuous segments; however, we found breaks in the ROH when plotted across these regions of elevated heterozygosity (Figure 3). Using BLAST (Agarwala et al., 2016), we were able to detect functional genes within eight of these islands of heterozygosity with various functions including signaling regulation, nucleotide binding, and the regulation of feeding behaviour. Several of the genes we identified have functions associated with immunity (Figure 3A-D), including: tyrosine kinase (TxK), a member of the PRAME family, and an immunoglobulin superfamily member (IgSF10). An island of heterozygosity on Scaffold 2797 (Figure 3D) contains prolactin (PRL), a gene best known for its role in reproduction, which has also been associated with immunity (Borba, Zandman-Goddard, & Shoenfeld, 2018). Some of the genes we detected have known polymorphisms, such as UDP-Glucuronosyltransferase (UGT) on Scaffold 3054 (Figure 3E), which functions in the elimination of toxins (Miners, McKinnon, & Mackenzie, 2002). We also found heterozygosity had been maintained within a region on Scaffold 3761 (Figure 3F) that was identified as Ankyrin Repeat Domain 26 (ANKRD26), a gene associated with feeding behaviour (Bera et al., 2008). When the caribou scaffolds are aligned to the bovine genome (National Center for Biotechnology Information, 2016), the scaffolds map to the respective bovine chromosomes that include each of these genes, validating our findings.

**FIGURE 3.**
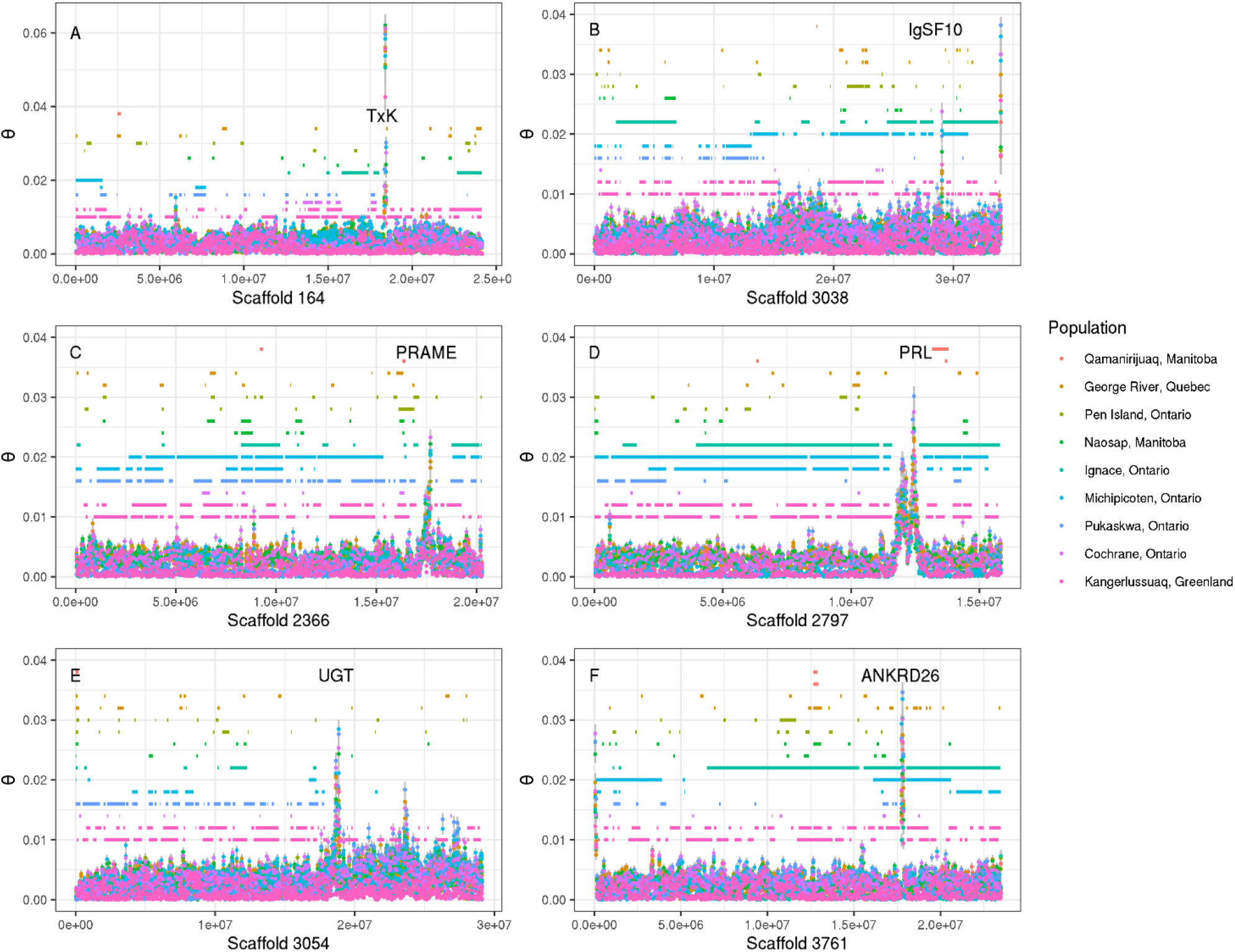
Heterozygosity estimates and Runs of Homozygosity (ROH) across select scaffolds of the caribou genome. Points represent local heterozygosity estimates calculated as Waterson’s θ every 50kb for each individual, with standard error bars. Horizontal line segements represent locations of ROH predicted by a hidden markov model for each individual. Colours represent caribou populations. Annotated labels on plots refer to genes identified within islands of heterzygosity. Note the Y-axis of plot A is 0.06; the Y-axes of plots B-F is 0.04.

## 4 DISCUSSION

We analyzed high-coverage whole genomes to investigate inbreeding extent in caribou from divergent populations in Canada and Greenland. We quantified ROH in high-coverage whole genomes of boreal caribou from small, isolated populations in the southern caribou range of Ontario, Canada, in comparison to caribou from the continuous range of Ontario, other caribou ecotypes in Canada, and western Greenland; populations presenting divergent evolutionary histories, differing in population size and extent of isolation. As predicted, we found boreal caribou from the southern range of Ontario had a relatively high proportion of their genomes in ROH (F_ROH_ = 0.1000, 0.2398), where recent range contraction has resulted in small and isolated populations (Figure 1). We also observed the longest average ROH in this region, namely in boreal caribou from Michipicoten Island, and Ignace Ontario, confirming their inbreeding occurred recently. Conversely, we found the lowest levels of inbreeding in barren-ground caribou from the Qamanirijuaq population in Manitoba, which was predicted as this large population has not experienced dramatic historic or recent declines (COSEWIC, 2016). All of the other Canadian caribou populations investigated had relatively low levels of inbreeding (Table 1). We examined two caribou from the Kangerlussuaq population in western Greenland, as previous research has indicated inbreeding in this population (Jepsen et al., 2002; Taylor et al., 2020), and found an extremely high proportion of ROH (F_ROH_ = 0.3373, 0.5017). Their genomes were comprised of some long ROH, although they were shorter than expected based on F_ROH_ (Figure 2), indicating their inbreeding has occurred over a longer period of time than that of other populations, as we predicted. Finally, we predicted that we would find genes associated with deleterious recessive alleles or heterozygote advantage within regions where balancing selection has maintained islands of heterozygosity despite inbreeding. We detected islands of heterozygosity on 18 of the 40 scaffolds examined and found breaks in the ROH when plotted across the islands. We identified functional genes within several of the islands that are associated with various functions, including immunity, the elimination of toxins, and a deleterious recessive condition associated with the regulation of feeding behaviour.

### 4.1 Inbreeding histories

We observed the lowest inbreeding estimates in barren-ground caribou from the Qamanirijuaq population in Manitoba. These caribou occur in large populations on the tundra and previous research has indicated the barren-ground caribou included in this study have admixed with other caribou ecotypes (Taylor et al., 2020), which was reflected by their low inbreeding estimates; in fact, the only negative inbreeding coefficients (F) observed were in barren-ground caribou from the Qamanirijuaq population, indicating outbreeding has occurred.

We found the highest prevalence of genomic inbreeding in caribou from western Greenland, whose genomes were 0.3373 and 0.5017 in ROH. Although extreme, these results are comparable to the uppermost ROH estimates of other studies; for example, inbred Scandinavian-born wolves have up to 0.54 of their genome in ROH (Kardos et al., 2018). The two individuals from western Greenland also shared a large proportion of those ROH that were IBD (92%). This is notably higher than other values in the literature. For example, a recent study of pumas reported a maximum of 36% IBD between two individuals who had similar inbreeding estimates as the western Greenland caribou (F_ROH_ = 0.5 – 0.6). However, our results also show that the two caribou from Greenland were 2^nd^ or 3^rd^ degree relatives, which elevates the proportion of the genome that is shared, or IBD.

In recent decades, native caribou of western Greenland have experienced population reductions of up to 90% (Jepsen et al., 2002). A microsatellite study of several caribou regions in western Greenland investigated the region included in our study, near Kangerlussuaq, and found high inbreeding coefficients at two of five loci investigated (Region 1; Jepsen et al., 2002). Despite evident inbreeding, caribou from the Kangerlussuaq region were polymorphic at all 5 loci investigated, unlike another isolated region that had lost several polymorphisms (Jepsen et al. 2002). Our analysis of whole genome sequences echoes this finding, as we find islands of heterozygosity maintained within several genes, despite extremely high inbreeding estimates (F_ROH_ = 0.3373, 0.5017). The high heterozygosity within these genes suggests they may be under balancing selection; thus, loss of heterozygosity at these loci may result in lowered fitness, or inbreeding depression. Although we also observed long ROH in caribou from western Greenland, they were shorter than predicted based on FROH, suggesting inbreeding occurred more historically in western Greenland than it has in the other inbred populations investigated. Our results indicate western Greenland caribou have undergone inbreeding over a longer time scale, which may have allowed for the purging of deleterious alleles. The maintenance of islands of heterozygosity within ROH suggests that inbred caribou may have enough genomic variation to avoid the fitness consequences of inbreeding depression. Indeed, despite our finding that caribou from the Kangerlussuaq region of western Greenland are extremely inbred, previous studies have found this population and the neighbouring population in Akia have notably high fertility, and are two of the only caribou populations in the world where twinning has been observed (Cuyler & Østergaard, 2005).

We examined the lengths of ROH to estimate if inbreeding occurred recently, resulting in long, continuous ROH, or historically, resulting in many short ROH. The longest average ROH were observed near the current southern edge of the boreal caribou range in Ignace, Ontario, suggesting recent isolation caused by range contraction. This result is congruent with the findings of a microsatellite study that suggested recent genetic erosion, a decrease in connectivity, and an increase in inbreeding along the southern continuous range edge of boreal caribou in Ontario and Manitoba (Thompson, Klütsch, Manseau, & Wilson, 2019). The second longest average ROH was observed on Michipicoten Island, Ontario, an isolated population that has experienced several recent bottlenecks (Bergerud et al., 2007; Fletcher, 2017). We also found the two caribou sampled from Michipicoten Island were related to each other but were not related to a caribou from the nearby coastal range in Pukaskwa National Park (Table S1). Notably, the caribou from Cochrane has undergone less inbreeding than the other Ontario boreal caribou sampled and had the least IBD ROH with the Michipicoten Island caribou. Previous research based on microsatellite data suggested Michipicoten Island and Pukaskwa National Park belong to a different genetic cluster than the nearby boreal caribou from Cochrane (Drake et al., 2018). Additionally, a recent genomic study found that boreal caribou from Cochrane, Ontario are genetically more similar to eastern migratory caribou from Quebec than they are to boreal caribou from Ignace, Ontario (Taylor et al., 2020), providing further evidence that there may be divergent evolutionary histories between these populations.

### 4.2 Islands of heterozygosity

Despite vastly different inbreeding histories among populations, we found regions of high heterozygosity that were maintained across all individuals, regardless of population size or ecotype. We identified functional genes within each of these peaks, with various functions including signaling regulation, nucleotide binding, and the elimination of toxins. Several of the genes we identified are associated with immunity, such as prolactin (PRL), which has been linked to both reproduction and immunity (Borba et al., 2018), and tyrosine kinase (TxK), which plays a role in T-cell development (Sommers et al., 1999). Benazzo et al. (2017) found similar evidence of balancing selection for genes associated with immune and olfactory systems, where non-random peaks of variation were maintained despite extreme inbreeding in endangered brown bears. We also found a gene that may have maintained variation due to the presence of a deleterious recessive allele. We found high heterozygosity within a region that contains ANKRD26, a gene associated with the regulation of feeding behaviour. Previous research has demonstrated homozygosity at this gene results in metabolic defects in mammals, including increased obesity and insulin resistance (Bera et al., 2008).

In a recent study of two eastern migratory caribou populations, correlations between heterozygosity and fitness (HFCs) found no evidence of inbreeding depression (Gagnon et al., 2019). Notably, eastern migratory caribou from one of these population were also included in our study and had relatively low inbreeding estimates (F_ROH_ = 0.05-0.07). The inbreeding levels in that population may be too low to trigger inbreeding depression, but it is also possible that the effects of inbreeding depression have been resisted by selection. Balancing selection can prevent the unmasking of deleterious recessive alleles or maintain heterozygote advantage thereby preventing the expression of inbreeding depression (Hedrick & Garcia-Dorado, 2016). In this study, we found islands of heterozygosity at specific gene regions across populations differing in population size and extent of isolation. This result suggests that strong balancing selection could be maintaining heterozygosity even in the face of extreme inbreeding. Balancing selection, specifically negative-frequency dependent selection, has also recently been suggested to be a mechanism maintaining phenotypic polymorphisms in caribou along an environmental gradient in western Canada (Cavedon et al., 2019).

### 4.3 Conclusions

We used runs of homozygosity (ROH) to quantify inbreeding in caribou from populations representing divergent evolutionary histories, differing in population size and extent of isolation. We explored the extent of inbreeding and the maintenance of genomic variation in high-coverage whole genomes of boreal caribou from small, isolated populations in the southern caribou range of Ontario, Canada, in comparison to caribou from the continuous range of Ontario, other caribou ecotypes in Canada, and western Greenland. We found divergent demographic histories among populations, particularly within the boreal ecotype, where we found low levels of inbreeding in caribou from the continuous boreal range, and elevated inbreeding estimates in populations that have become isolated due to recent range contraction. We also observed the longest average ROH in these isolated boreal caribou populations, confirming their inbreeding occurred recently. We observed the greatest amounts of inbreeding in caribou from western Greenland, whose genomes were approximately a third to a half in ROH; although their ROH were shorter than expected, implying inbreeding has occurred over a longer time in western Greenland than it has in Ontario.

Across populations with divergent demographic histories, we infer the maintenance of variation within genes associated with various functions, including immunity, the removal of toxins, and a deleterious recessive condition, suggesting balancing selection is occurring despite extreme inbreeding. The maintenance of heterozygosity in these key regions may help resist the effects of inbreeding depression. To further investigate how inbreeding depression may be affecting these populations, researchers should examine beyond the islands of heterozygosity and attempt to identify genes located within ROH, especially those that are likely to be associated with heterozygote advantage or a deleterious recessive allele.

## ACKNOWLEDGEMENTS

Funding was provided through an NSERC Collaborative Research & Development (CRD) grant to P.J.W., NSERC CREATE and Discovery grants to J.B. and P.J.W., and a Garfield Weston Foundation Fellowship to K.S., Ontario Graduate Scholarships, and Ontario Species at Risk Stewardship Program funding to K.S. The results reported are reflective of the independent analyses conducted for this study and do not necessarily reflect the views of the Government of Ontario. We are grateful to Dr. Art Rodgers for his assistance with sample collection and advice on this paper. We thank Bridget Redquest and Austin Thompson for extracting DNA, and The Centre for Applied Genomics (TCAG) at the Hospital for Sick Children (Toronto, Ontario) for library preparation and sequencing. We also acknowledge the facilities of the Shared Hierarchical Academic Research Computing Network (SHARCNET: www.sharcnet.ca) and Compute Canada/ Calcul Canada gme-665-ab. Compute Canada (RRG gme-665-ab), and Amazon Cloud Computing for high-performance computing services. We are grateful for the efforts of those who provided tissue samples including the Government of Manitoba, Government of Ontario, Government of Greenland, and Environment and Climate Change Canada.

## DATA ACCESSIBILITY

The raw reads of 12 individuals are available at the National Centre for Biotechnology (NCBI) under the BioProject Accession no. PRJNA 634908. The raw reads of the remaining three individuals will be made available by time of publication.

## AUTHOR CONTRIBUTIONS

K.S., J.B., P.J.W., and M.M. conceived and designed the study. J.B., P.J.W., and M.M. oversaw the research as Co-Principal Investigators. K.S. and S.K. performed bioinformatic analyses with guidance from R.S.T. and R.L.H. K.S. wrote the manuscript and J.B., R.S.T, M.M., and P.J.W. provided feedback and edited the manuscript.

