## Supplementary Materials for "Genomic islands of heterozygosity maintained across caribou populations despite inbreeding"

MOLECULAR ECOLOGY

### Table of Contents:


| | |
|---|---|
| **Table S1** | Page 2 |
| **Table S2** | Page 3 |
| **Figure S1** | Page 4 |
| **Figure S2** | Page 5 |
| **Figure S3** | Page 6 |
| **Figure S4** | Page 6 |

MOLECULAR ECOLOGY

**Table S1.** Relatedness matrix between individual caribou based on the KING inference of relatedness, calculated from a joint genotyped VCF file containing 28 246 751 SNPs. Column and row headers indicate individual ID number.

| **Sample ID** | 21332 | 21350 | 27689 | 27694 | 34590 | 20917 | 35324 | 35326 | 39590 | 39650 | 39651 | 39653 | 39654 | 41660 | 41667 |
|---|---|---|---|---|---|---|---|---|---|---|---|---|---|---|---|
| 21332 | 0.5 | 0.0383 | -0.0836 | -0.0716 | -0.0577 | -0.0469 | -0.0657 | -0.0438 | -0.1867 | -0.1684 | -0.1650 | -0.2910 | -0.0821 | -0.3521 | -0.3561 |
| 21350 | - | 0.5 | -0.0836 | -0.0698 | -0.0559 | -0.0473 | -0.0640 | -0.0422 | -0.1855 | -0.1681 | -0.1635 | -0.2896 | -0.0804 | -0.3530 | -0.3569 |
| 27689 | - | - | 0.5 | 0.0347 | -0.0242 | -0.0208 | -0.0518 | -0.0425 | -0.1412 | -0.1245 | -0.1218 | -0.2444 | -0.0091 | -0.6140 | -0.6188 |
| 27694 | - | - | - | 0.5 | -0.0122 | -0.0057 | -0.0356 | -0.0302 | -0.1227 | -0.1052 | -0.1023 | -0.2251 | 0.0043 | -0.5831 | -0.5878 |
| 34590 | - | - | - | - | 0.5 | 0.0408 | -0.0012 | 0.0072 | -0.0884 | -0.0717 | -0.0701 | -0.1894 | 0.0058 | -0.5789 | -0.5838 |
| 20917 | - | - | - | - | - | 0.5 | 0.0044 | 0.0146 | -0.0840 | -0.0674 | -0.0644 | -0.1846 | 0.0130 | -0.5530 | -0.5587 |
| 35324 | - | - | - | - | - | - | 0.5 | -0.0038 | -0.1236 | -0.1047 | -0.1002 | -0.2276 | -0.0290 | -0.5812 | -0.5874 |
| 35326 | - | - | - | - | - | - | - | 0.5 | -0.1195 | -0.1008 | -0.0977 | -0.2224 | -0.0162 | -0.5493 | -0.5536 |
| 39590 | - | - | - | - | - | - | - | - | 0.5 | -0.0315 | -0.0450 | -0.3182 | -0.0996 | -0.8624 | -0.8716 |
| 39650 | - | - | - | - | - | - | - | - | - | 0.5 | 0.0772 | -0.2321 | -0.0824 | -0.8234 | -0.8299 |
| 39651 | - | - | - | - | - | - | - | - | - | - | 0.5 | -0.2270 | -0.0810 | -0.8129 | -0.8206 |
| 39653 | - | - | - | - | - | - | - | - | - | - | - | 0.5 | -0.2019 | -1.1274 | -1.1400 |
| 39654 | - | - | - | - | - | - | - | - | - | - | - | - | 0.5 | -0.6248 | -0.6298 |
| 41660 | - | - | - | - | - | - | - | - | - | - | - | - | - | 0.5 | 0.0930 |
| 41667 | - | - | - | - | - | - | - | - | - | - | - | - | - | - | 0.5 |

MOLECULAR ECOLOGY

**Table S2.** Scaffolds where global heterozygosity (θ) exceeded 0.02, selected from the 40 largest scaffolds of the caribou genome. Column headers indicate individual ID number of each caribou. Y or N (yes or no) indicates whether heterozygosity exceeded 0.02 for each individual. The gene column indicates genes that we were able to identify within each region using BLAST.

| Scaffold | 21332 | 21350 | 27689 | 27694 | 20917 | 34590 | 35324 | 35326 | 39590 | 39650 | 39651 | 39653 | 39654 | 41660 | 41667 | Gene |
|---|---|---|---|---|---|---|---|---|---|---|---|---|---|---|---|---|
| 164 | Y | Y | Y | Y | Y | Y | Y | Y | Y | Y | Y | Y | Y | Y | Y | TxK |
| 1010 | N | N | N | N | N | N | N | Y | N | Y | Y | N | Y | N | N | |
| 1653 | N | N | N | N | N | N | N | N | Y | Y | N | Y | Y | N | N | |
| 1870 | N | N | N | N | N | N | N | Y | N | Y | N | Y | N | N | N | |
| 1991 | Y | Y | Y | Y | Y | Y | Y | Y | Y | Y | Y | Y | Y | Y | N | |
| 2215 | Y | Y | Y | Y | Y | Y | Y | Y | Y | Y | Y | Y | Y | Y | Y | |
| 2246 | Y | Y | Y | Y | Y | Y | Y | Y | Y | Y | Y | Y | Y | Y | Y | |
| 2249 | Y | Y | Y | N | Y | Y | N | Y | N | Y | Y | Y | N | N | N | |
| 2366 | Y | N | N | N | N | N | N | Y | N | N | N | N | Y | N | N | PRAME |
| 2797 | Y | N | Y | Y | Y | N | N | Y | N | N | Y | Y | Y | N | N | PRL |
| 2948 | Y | Y | Y | Y | Y | Y | Y | Y | Y | Y | Y | Y | Y | Y | Y | multiple |
| 3038 | N | N | Y | Y | Y | N | N | Y | N | Y | Y | Y | Y | Y | N | IGSF10 |
| 3054 | N | N | N | N | N | N | N | Y | N | N | Y | Y | Y | N | N | UGT |
| 3160 | Y | Y | N | Y | N | N | N | Y | N | Y | Y | Y | Y | Y | N | |
| 3284 | Y | Y | Y | Y | Y | Y | Y | Y | Y | Y | Y | Y | Y | N | N | RWDD1, TAAR |
| 3761 | Y | Y | Y | Y | Y | N | N | Y | N | Y | Y | Y | Y | Y | N | ANKRD26 |
| 3936 | Y | Y | N | Y | Y | Y | N | Y | Y | Y | Y | Y | Y | Y | N | |

MOLECULAR ECOLOGY

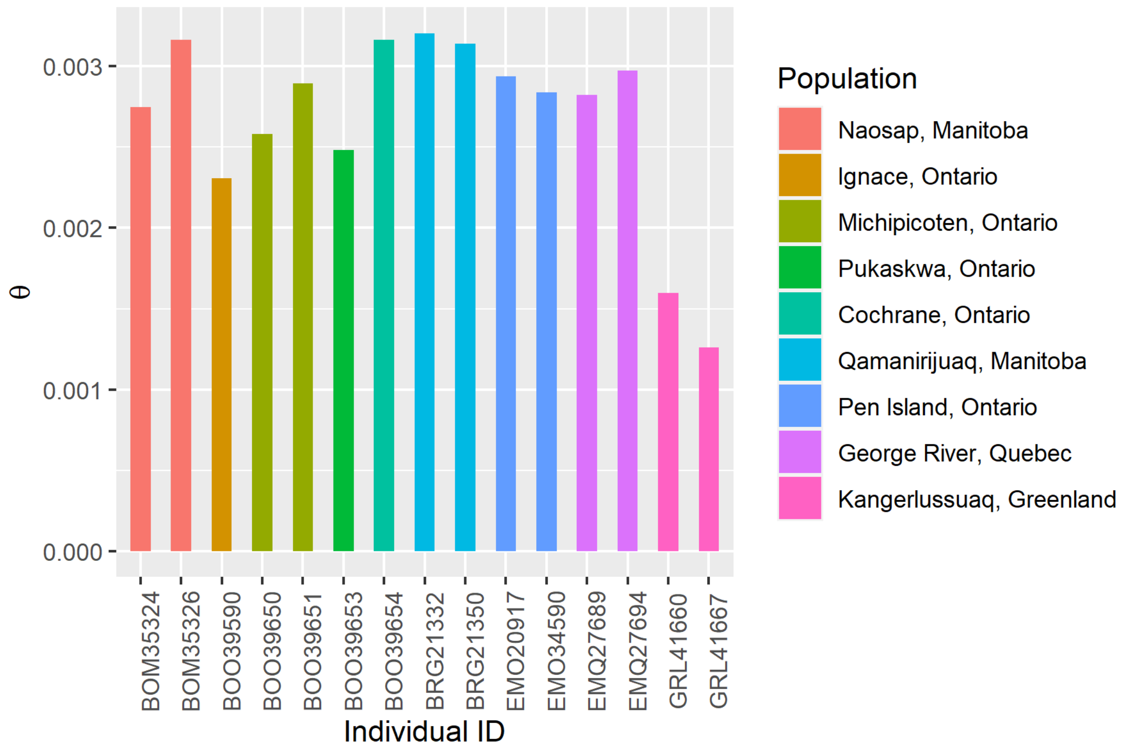


**Figure S1**. Average genome-wide heterozygosity (Watterson's θ) for each caribou calculated from individual high-coverage BAM files.

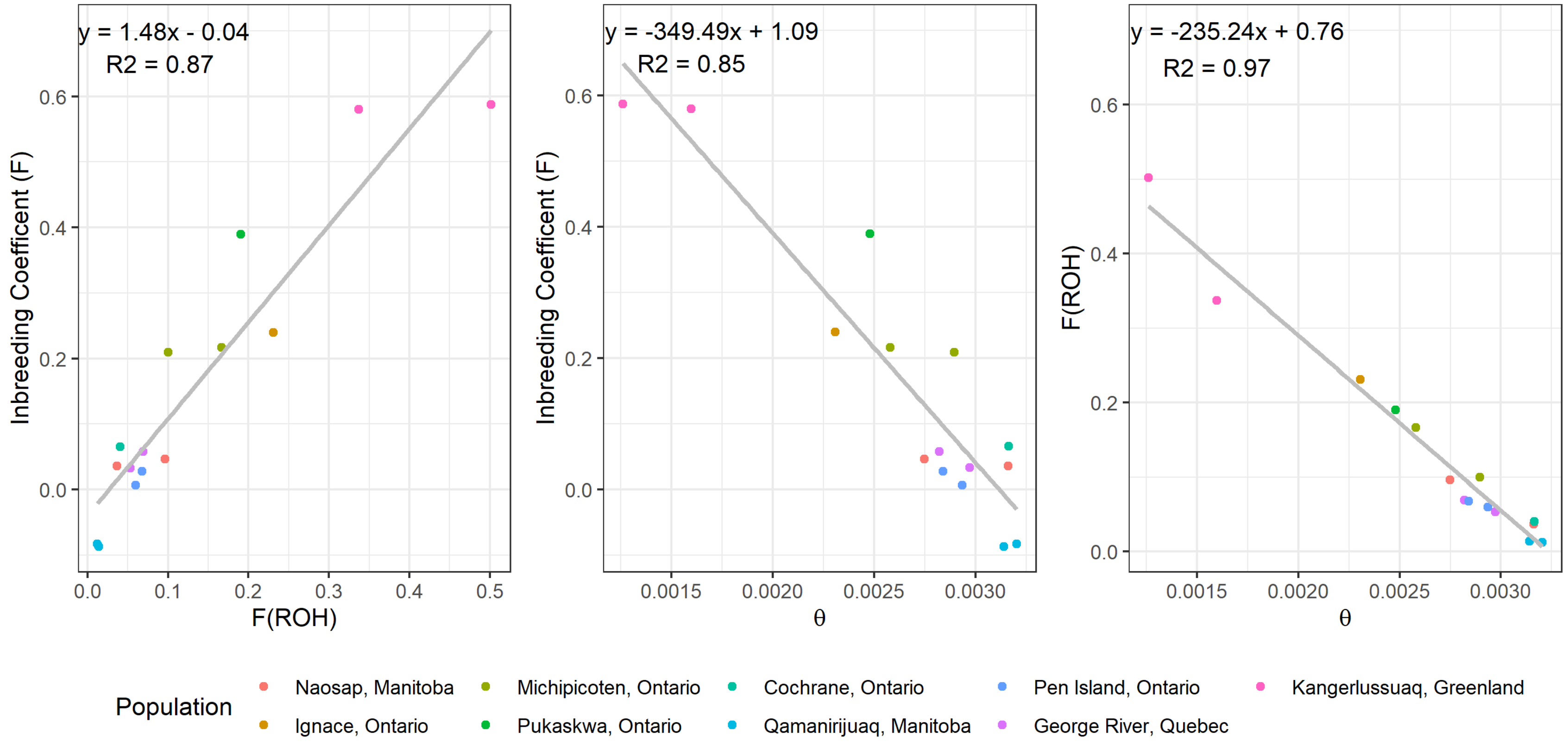


**Figure S2.** Linear model correlations between inbreeding estimates. The inbreeding coefficient (F) was calculated from a joint VCF file containing 28 246 751 SNPs, $F_{(ROH)}$ and genome wide-heterozygosity (Watterson's θ) were calculated from individual high-coverage BAM files. The equation reflects the line of best fit and $R^2$ is the adjusted R-squared value. All correlations were significant ($p < 0.0001$).

MOLECULAR ECOLOGY

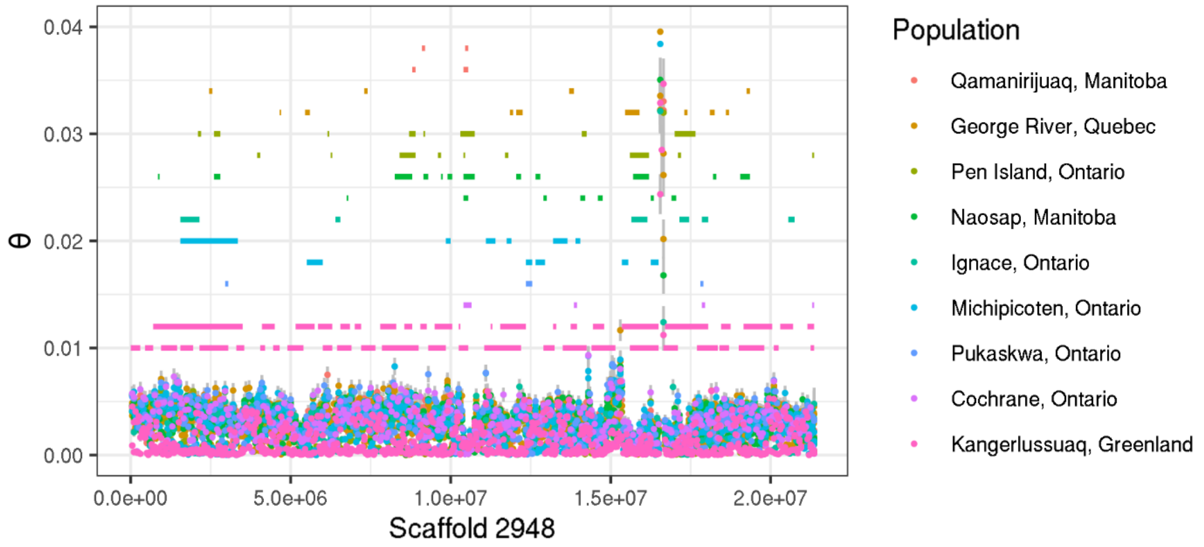


**Figure S3.** Points represent local heterozygosity across Scaffold 2948, estimated as Waterson's θ with standard error bars for each caribou. Horizontal lines represent ROH segments predicted by a hidden markov model. Colours represent populations caribou were sampled from. We identified multiple genes around the island of heterozygosity, including AFG3L2, GNAL, CHMP1B, and CCP110. All of the genes identified are located on the 24$^{th}$ chromsome of the bovine genome, which Scaffold 2948 aligns to.

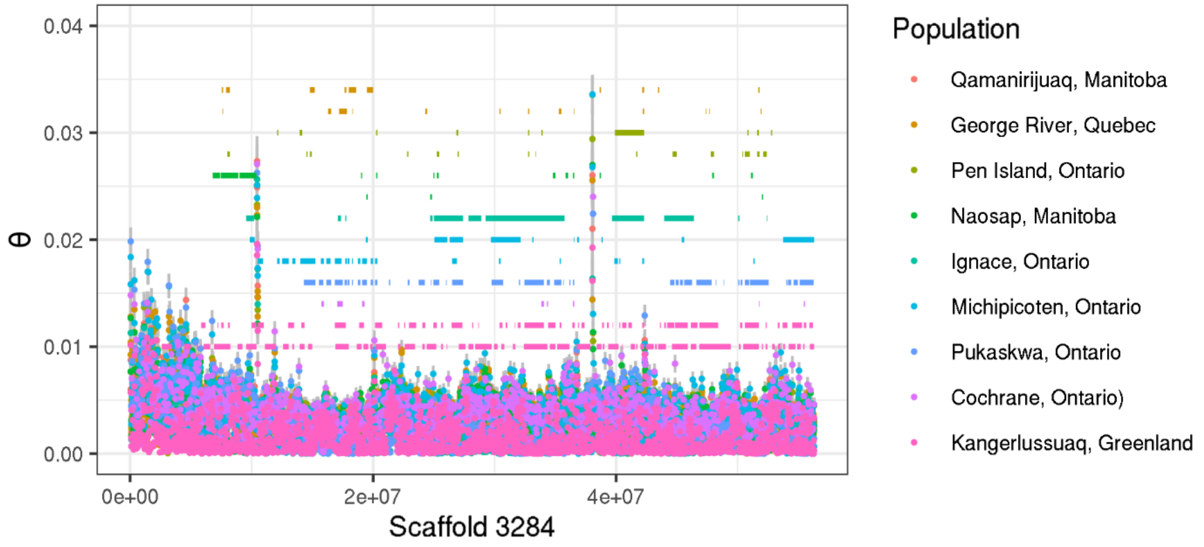


**Figure S4.** Points represent local heterozygosity across Scaffold 3284, estimated as Waterson's θ with standar error bars for each caribou. Horizontal lines represent ROH segments predicted by a hidden markov model. Colours represent populations caribou were sampled from. We identified the gene RWDD1 within the first island of heterozygosity, and TAAR within the second island. Both genes are located on the 9$^{th}$ chrosomome of the bovine genome, which Scaffold 3284 aligns to.
